# Association of *Blastocystis* and Gut Microbiota in Type 2 Diabetic Mellitus Patients and non-Diabetic Individuals

**DOI:** 10.1101/2024.01.29.577889

**Authors:** Nurul Saadah Mohd Shaari, Wan Shahida Wan Sulaiman, Mohd Rahman Omar, Nadeeya’Ayn Umaisara, Ii Li Lee, Tengku Shahrul Anuar, Noradilah Samseh Abdullah

**Author notes:** These authors contributed equally to this work (Nurul Saadah Mohd Shaari, Wan Shahida Wan Sulaiman, Mohd Rahman Omar, Nadeeya’Ayn Umaisara, Noradilah Samseh Abdullah). These authors also contributed equally to this work.

## Abstract

The influence of anaerobic protozoan *Blastocystis* on human gut health is not well understood. While *Blastocystis* species frequently inhabit the gut, their clinical importance and ecological function remain ambiguous. A study on *Blastocystis* was carried out enrolling a total of 203 participants including T2DM patients and non-diabetic individuals to evaluate the prevalence of *Blastocystis* and its association in gut microbiota. *Blastocystis* subtypes were identified by PCR and faecal microbiome was accessed by targeting V4 region of the bacterial 16S ribosomal gene. The prevalence of *Blastocystis* in T2DM was 25.49% and 17.82% in non-diabetic individuals with the most prevalent subtype on total population was ST3, followed by ST1 and ST2. The composition of gut microbiota was significantly different between *Blastocystis*-positive and *Blastocystis*-negative individuals. *Blastocystis* carriage was positively associated with higher alpha diversity in T2DM patients and non-diabetic individuals. Interestingly, at the phylum level, the T2DM group had an obvious increase of Bacteroidetes and a marked increase of Actinobacteria with the present of *Blastocystis*. The findings suggested that the presence of *Blastocystis* was linked to increased diversity and richness in the gut bacterial composition, signifying at a potentially beneficial association between *Blastocystis* and the gut microbiota.

**Author Summary:** Type 2 Diabetes Mellitus Patients (T2DM), a prevalent global disease, affects a significant portion of the population across the world. Thus, there is need to better understanding on *Blastocystis* infection among T2DM that could lead to the alteration toward gut health. We evaluated the association between *Blastocystis* and gut microbiota, where involving two groups; T2DM patients and non-Diabetic individuals. The research revealed a higher *Blastocystis* in T2DM patients compared to non-diabetic individuals, emphasizing on assumption toward its pathogenicity. However, amplicon-based sequencing of 16S rRNA genes indicates that *Blastocystis* carriers exhibit increased gut microbiota diversity. Our result suggested that, *Blastocystis* highlighted its potential role as a component of a balanced microbiota. Notably, optimal alteration in Actinobacteria and Bacteroidetes may contribute to the several gut health. Hence, the study could prompt for further exploration regarding of *Blastocystis* subtypes and gut microbiota specifically in T2DM to propose for more precise assessment of *Blastocystis* and gut microbial diversity.

## Introduction

Anaerobic protozoan *Blastocystis* is one of the most prevalent intestinal parasites in humans and has been controversial for decades. Currently, analysis of the ribosomal RNA gene (SSU rDNA) revealed up to 33 genetically distinct strains capable of colonising a variety of hosts, including humans and animals (1,2). In Malaysia, the mean prevalence of *Blastocystis* in humans reported to be around 19.25%, with a range of 9.17-40.30%. According to *Blastocystis* subtype distribution, humans have been reported to have ST1-4 and ST6 subtypes, with ST3 (52.51%) being the predominant subtype. There are five common subtypes: ST1 (27.16%), ST2 (8.45%), ST6 (5.43%), ST7 (3.42%), and ST4 (1.2%) (3,4).

Previous studies has suggested a potential link between *Blastocystis* and type-2 diabetes mellitus (T2DM) based on the basic prevalence data (5,6). In T2DM, insulin resistance does not solely result from being overweight but also involves a complex interplay of multiple factors such as gut ecosystem and immune response, where they might potentially influence by the pathogenesis of T2DM (7). Recent studies demonstrated that the diversity of gut microbiota has significantly declined in T2DM comparing to healthy control (8,9). However, the infection by *Blastocystis* in diabetic patients associated with gut microbiota is not completely discovered.

Gut microbiota comprises thousands of diverse microbes including bacteria with the major bacteria species found in the human gut, which make up more than 90% of the gut microbiota belong to one or two bacteria phyla: Firmicutes and Bacteroidetes (10). Gut microbiota is essential to the host’s physiological activities; altering the delicate host-microbiota interaction equilibrium may affect the onset underlying various metabolic disorders, including T2DM (11). Dysbiosis will occur if the microbiota is imbalance (12). The large number of previous studies indicated that the presence of *Blastocystis* would increase the diversity of gut microbiota, and healthier people frequently have higher gut microbiota diversity. The parasite could be considered as a commensal of healthy gut health (13,14). However, there contradictory findings that suggested *Blastocystis* may be linked to dysbiosis (15–17). Recent research by Defaye and colleagues has shown that alterations in the microbiota that are linked to visceral hypersensitivity are caused by *Blastocystis* infection in rats (15). *Oscillospira*’s relative abundance increased whereas *Clostridium*’s relative abundance decreased as a result of these alterations.

Although many studies have found a bilateral correlation between the presence of *Blastocystis* and the composition of the gut microbiota, it is unclear whether the gut conditions and gut microbiota perturbation lead to higher *Blastocystis* colonisation in the gut or whether *Blastocystis* colonisation may lead to dysbiosis (18). Facing these contradictory results, the need for additional data for microbiota modification associated with *Blastocystis* colonization should be performed. Therefore, the present study was conducted to evaluate the prevalence of *Blastocystis* and its association with gut microbiota in T2DM patients and non-diabetic individuals.

## Methods

### Patient recruitment and sample collection

This was a cross section study with a non-probability sampling conducted between February 2021 to November 2022. One hundred and two T2DM patients and 101 non-diabetic participants ranging from men and women above 18 years old consented to this study. Diabetic patients who were regularly attending Endocrine Clinic and Primary Medical Centre, Universiti Kebangsaan Malaysia and fulfilled the inclusive criteria were recruited for this study. Non-diabetic participants were recruited from volunteers who did not have any serious disease and fulfil the exclusion criteria for diabetic group. They were approached during their daily medical check-up; some were from the community living around Klang Valley or areas that have similar sociodemographic as the diabetic group. Participants who consumed antibiotics and probiotics less than 3 months before stool collection were excluded from the study. This study was approved by USIM(USIM/JKEP/2020-82) and UKM(UKMPPI/111/8/JEP-2019-848) ethics committees.

### Faecal samples and DNA extraction

A collection of faecal samples from subjects was submitted to the laboratory by guidelines. The samples were transferred immediately into several tubes and stored at -80 °C until processing. Then, the genomic DNA (gDNA) was directly extracted from 203 unfixed stool samples using QIAamp® PowerStool Pro DNA Kit (Qiagen, Germany) according to the manufacturer’s protocol.

### Genus and *Blastocystis* subtypes determination

The genomic DNA from faeces from all subjects were subjected to polymerase chain reaction (PCR) analyses to detect the presence of *Blastocystis* by using barcoding primer with the primers RD5 (5′-ATC TGG TTG ATC CTG CCA GT-3′), and BhRDr (3′-GAG CTT TTT AAC TGC AAC AACG-5′) amplifying a ∼ 600-base pair (bp) fragment of the 1.8 kbp small subunit ribosomal RNA (SSU rRNA) gene. This region of the SSU-rRNA gene has shown to provide sufficient information for differentiating subtypes of *Blastocystis* (19). 2 µL of extracted DNA was added to an amplification mixture containing 1 µl (10 µM) of primers and 12.5 μl of PCR Master Mix Plus (Promega, USA). To each mixture, nuclease-free water was added to provide a final reaction volume of 25 μl. Nuclease-free water was used as a negative control, and *Blastocystis* ST3 DNA, obtained from the stool samples of colonized volunteers, was used as a positive control. The amplification profile consisted of 30 cycles of denaturation, annealing and extension at 95 °C, 63 °C and 72 °C respectively, with a final extension step of 4 minutes at 72 °C. PCR products (5 μl) were analysed on 1.5% agarose gel using electrophoresis and visualized under UV light. The sequences were analysed by the BLAST website (https://blast.ncbi.nlm.nih.gov/Blast.cgi?PAGE_TYPE=BlastSearch). STs and ST (SSU-rDNA) alleles were called using the sequence query facility in the *Blastocystis* Sequence Typing website available at https://pubmlst.org/bigsdb?db=pubmlst_blastocystis_seqdef&page=sequenceQuery. Subtypes were identified by determining the exact match or closest similarity to all known *Blastocystis* subtypes according to the classification by Stensvold (20).

### 16S rRNA-amplicon based sequencing

100 samples were subjected to bacterial diversity characterization by amplicon sequencing of the 16S rRNA gene using the Illumina MiSeq platform by NGS servicer (First base, Malaysia). 16S rRNA V4 Forward (5′-CCTACGGGNGGCWGCAG -3′) and 16S rRNA V4 -R (5′-GACTACHVGGGTATCTAATCC’3) were used because the V4 hypervariable region has been reported to be the most informative region for describing bacterial communities. For the sequencing process, microbial amplicon libraries were built and sequenced until a minimum expected raw depth of 300 thousand reads per sample was reached.

### Bioinformatic analysis

Sequence adapters and low-quality reads were removed from the paired end reads before the first 60 000 raw reads were extracted out using BBTools (DOE Joint Genome Institute). The forward and reverse reads were merged using QIIME. The error reads low-quality regions and chimaera errors were removed and corrected with the DADA2 pipeline (https://benjjneb.github.io/dada2/). The resulting data was in the form of an amplicon sequence variant (ASV) and proceeded with the taxonomic classification by using scikit-learn (https://scikit-learn.org/stable/) and Naive Bayes classifier against the database SILVA (release 132). Analysis was performed in R Studio version 3.6.2 by using phyloseq (https://www.bioconductor.org/packages/release/bioc/html/phyloseq.html). Alpha diversity metrics (richness and Shannon’s index) were calculated using the phyloseq R package based on rarefied OTU counts.

### Statistical analysis

A descriptive analysis of the variables studied was conducted. For all continuous values, normality assumptions were evaluated using the Kalmogorov-Smirnov and Shapiro Wilk. The quantitative variables were summarized in terms of means and standard deviation or median and interquartile range and the qualitative variables were summarized in frequencies and proportions. Alpha diversity metrics were compared by Mann–Whitney tests; for comparisons involving more than two groups, Bonferroni’s correction was applied. Probability (*p*) values p < 0.05 were considered to indicate statistically significant differences. The statistical analysis was performed using IBM SPSS Statistic 25.

## Results

### Prevalence of *Blastocystis* and subtypes

Based on PCR detection, the prevalence rate of *Blastocystis* infection in the diabetic group was 25.49% (26/102) while 17.82% (18/101) samples were detected in the *Blastocystis* infection in the non-diabetic group. *Blastocystis* infection was higher in diabetic patients compared to non-diabetic group. In diabetic patients, the detected subtypes were ST3 (34.61) ST1(26.9%), ST2 (15.38%), ST4 (11.54%), ST6 (7.69%) and ST7 (7.69%). Meanwhile, in non-diabetic individuals, it was found that ST3 (33.3%) was the most common subtype detected followed by ST1 (27.7%), ST2 (22.2%), ST4 (11.1%) and ST6 (5.6%).

### Association of *Blastocystis* colonization with richness and diversity of gut microbiota

Figure Analysis on the 16 rRNA sequence data set by revealing a significant decrease in bacterial richness (Observed ASV, Chao 1 and Fisher) in T2DM patients compared to the non-diabetic group (Figure 1). Diversity (Shannon function and Simpson’s index) was also significantly different between T2DM patients and non-diabetic group. Interestingly, both bacterial richness and diversity increase in *Blastocystis* carriers in both T2DM and non-diabetic groups compared to negative *Blastocystis* participants.

**Figure 1:**
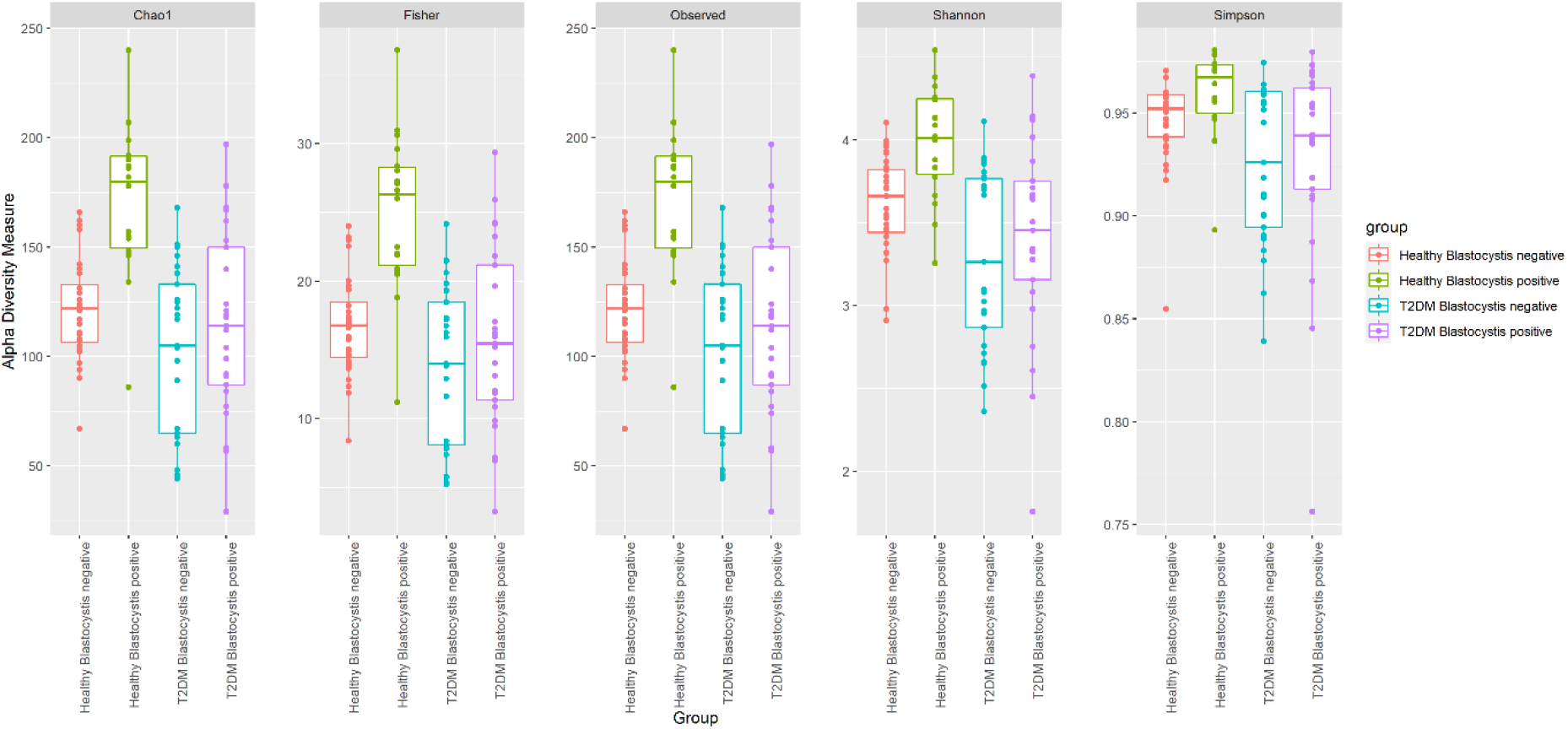
Alpha diversity measure differences between *Blastocystis* positive and *Blastocystis* negative faecal DNA samples in both T2DM and non-diabetic groups. Bacterial richness (Observed ASV, Chao 1 and Fisher) and bacterial diversity (Shannon’s diversity and Simpson’s index). All measures differed significantly (p < 0.05).

### Gut microbiome composition according to Blastocystis Colonization

The bacterial composition of the faecal sample from *Blastocystis* positive and *Blastocystis* negative patients were further analysed by measuring their sequencing read in the ASV table. At the phylum level, Figure 2 shows that several phyla were detected within the samples. The top abundance phyla; Actinobacteria, Proteobacteria, Bacteroidetes, Firmicutes and Verrucomicrobia in Table 1. Based on Table 1, Kruskal-Wallis analysis of the groups found that there were significant differences in the Actinobacteria and Bacteroidetes within four groups (p < 0.05). T2DM *Blastocystis* positive group had a higher abundance of Proteobacteria, Bacteroidetes and Firmicutes compared to T2DM *Blastocystis* negative group. However, there were no significant in the abundance of Proteobacteria, Firmicutes and Verrucomicrobia (p > 0.05).

**Table 1.** Abundance of bacterial phyla in T2DM and non-DM according to *Blastocystis* positive and *Blastocystis* negative group.

| Phyla | T2DM |  | Non-DM |  | p-value |
| --- | --- | --- | --- | --- | --- |
|  | *Blastocystis* | *Blastocystis* | *Blastocystis* | *Blastocystis* |  |
|  | positive | Negative | positive | Negative |  |
|  | (n=25) | (n=25) | (n=18) | (n=32) |  |
| Actinobacteria | 3839 (3826) | 5149(6535) | 1949 (2182) | 2603 (3353) | 0.003* |
| Proteobacteria | 910 (2780) | 588 (2597) | 1093(2240) | 645.50 (855) | 0.207 |
| Bacteroidetes | 4887(7438) | 1729 (5636) | 6539 (3265) | 6382 (4957) | p < 0.001* |
| Firmicutes | 13630 (5487) | 13525 (4313) | 14712.50 (2588) | 13418 (3769) | 0.974 |
| Verrucomicrobia | 0.00 (193) | 0.00(3) | 5.5 (67) | 0.00 (96) | 0.214 |

Data are expressed as median (interquartile range (IQR) for skewed data, analysed using the Kruskal-Wallis test for continuous variables in SPSS, p-value < 0.05 is significant.

**Figure 2.**
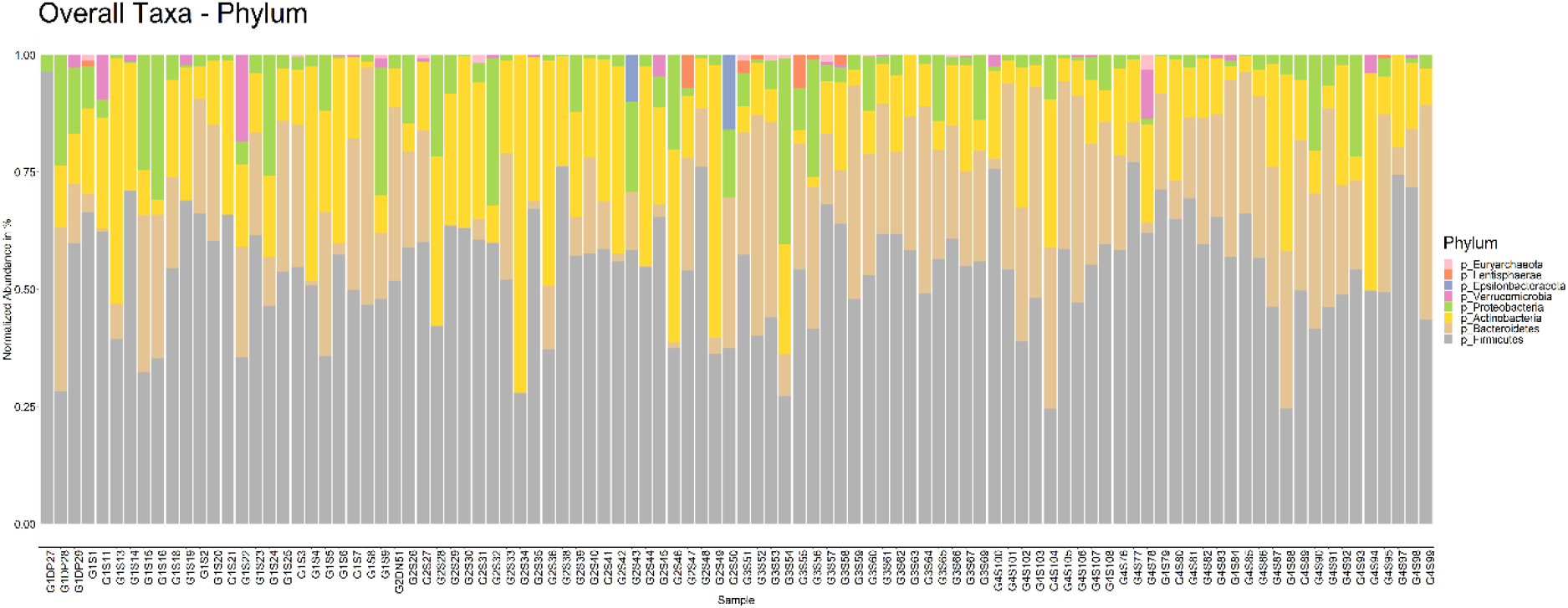
Bar chart of relative abundance at the phylum level between study participants.

## Discussion

Our study was motivated by previous literature where there were outgrowing numbers of research regarding the evaluation of gut microbiota in the presence of *Blastocystis* with the diverse findings on gut composition. The analysis of the intestinal microbiota of individuals with and without *Blastocystis* has piqued the scientific community’s interest because it may clarify the pathogenic potential, innocuousness, or even beneficial properties of this protozoan in humans (21). Recent studies showed that *Blastocystis* colonisation could be associated with discrete changes in gut microbiota composition especially in immunocompromised patients such as inflammatory bowel syndrome (IBS), inflammatory bowel disease (IBD) patients. The studies indicate that there might be a positive correlation between the presence of *Blastocystis* and changes in specific bacterial population. Specifically, a study conducted by Nourrisson and colleagues (22) has found that in male patients with *Blastocystis* colonization, there is an increase in *Lactobacilli* and decrease *Bifidobacterium* sp. and *F. prausnitzii*. This result suggests that *Blastocystis* colonization may lead to a decrease protective bacteria (22). According to previous studies, there was high prevalence of *Blastocystis* infection had been reported in T2DM compared to healthy individuals (5,6,23,24). Hence, our data represents the first study focusing on the diversity and abundance of gut composition concerning the presence of *Blastocystis* in both T2DM and non-diabetic Malaysian adults.

T2DM was described by patients particularly based on glucose level and consume conventional treatments and medication periodically or over the long term. Probiotics and antibiotic were exclusion criteria to avoid bias during the study of gut microbiota. Thus, patients enrolled in this study would be able to participate in comparable groups. In the present study, a high prevalence of *Blastocystis* infection detected in T2DM compared to healthy individuals, which is similar with previous studies that were conducted in Thailand, Iran and Brazil (5,6,24). T2DM in humans is linked to changes in the composition of the intestinal microbiota, which reduces the abundance of some universal butyrate-producing bacteria and increases the likelihood of various opportunistic pathogens such as *Blastocystis*. This parasite could facilitate colonization by other intestinal pathogens and changes in the intestinal microbiota diversity and composition (25,26).

Our study appeared to find a significant increase of alpha diversity in *Blastocystis* carriers from both T2DM and non-diabetic groups. Increasing in alpha diversity associated with *Blastocystis* carriage suggested that *Blastocystis* may be a component of healthy microbiota (27). Interestingly, most of the previous findings demonstrated that individuals colonized by *Blastocystis* typically have greater alpha diversity and various shifts in the composition of the gut microbiota than those who do not carry the organism (13, 27–29). The intestinal microbiota is highly variable among individual humans and its diversity is affected by factors like diet, socio-geographic setting, antibiotic use, disease, age and to a lesser degree, genetics (30). In conjunction with an alteration in the immunocompromised host, *Blastocystis* infection has been reported to diversify the gut microbiota in healthy individuals and those interactions may have downstream effects on host immunity and gut homeostasis (31–33). In a study by the Flemish Gut Flora Project (FGFP) on a western population cohort, a robust link was discovered between *Blastocystis* subtypes and microbiota community composition, surpassing host characteristic in explanatory strength. This association accounted for approximately 30% of *Blastocystis* prevalence in the non-clinical cohort, contrasting with 4% in Flemish IBD patients. The finding suggested that *Blastocystis* subtypes or even intra-subtypes within the prevalence genus were associated with human health (29,30,34). In a healthy gut, metabolites such as short-chain fatty acids are developed by fermentation by anaerobic bacteria and possibly by *Blastocystis* (35).

At the taxonomic phyla level, our results displayed that the highest relative abundant phyla were Firmicutes, Bacteroidetes, Proteobacteria and Actinobacteria for all the groups (Table 1), which agrees with previously reported findings. There was a significant difference in Actinobacteria and Bacteroidetes phyla within groups. The increase of Firmicutes had been observed in T2DM patients and non-diabetic individuals who were *Blastocystis* carrier. In addition, there was a reduced abundance of Bacteroidetes in diabetic individuals compared to healthy individuals. It was suggested that the change in both phyla in the T2DM group, a low microbial gene count and a dominance in genera *Bacteroides* and *Ruminococcus* was associated with a more remarkable ability to obtain energy from diet, systemic inflammation, adiposity, insulin resistance and dyslipidaemia (36). Moreover, Bacteroidetes are known to produce primarily acetate and propionate, whereas Firmicutes produce more butyrate, which has anti-inflammatory properties, regulates energy metabolism, and increases leptin levels (37,38). Despite not all phyla identifying statistically significant differences in the individuals colonized by *Blastocystis*, there was a probably lower abundance of Bacteroidetes and an increase in Proteobacteria. Similarly, previous study from school age children from Colombia demonstrated that phylum Firmicutes was the predominant taxonomic unit in both groups. In addition, there was a high proportion of Bacteroidetes in *Blastocystis-free* individuals while there was a higher relative abundance of the phylum Proteobacteria where the family *Enterobacteriaceae* shows a probable increase in abundance in children colonized by *Blastocystis*. Despite there were no statistical changes between the groups with and without *Blastocystis*, the result suggested a greater richness and diversity of the intestinal microbiota in the colonized individuals (39). As proposed by Gabrielle and Co-workers, ST3 was associated with healthy microbiota and may be related to a greater richness of intestinal microbiota (40). This finding also was supported by a study from Mexico where ST3 was described in the FASCA cohort and UNEME cohort with no association in the intestinal low Firmicutes/Bacteroidetes (F/B) ratio (13). It is interesting to note the presence of *Blastocystis* and *Prevotella copri*, which both strongly correlate with favourable glucose homeostasis and a reduction in the estimated visceral adipose tissue mass as markers of improved postprandial glucose response (41).

Significantly, his study concluded that the presence of *Blastocystis* is a sign of a balanced gut microbiota. Our findings imply that *Blastocystis* may be associated with an increase in richness because it may have a positive impact on the level of gut microbiota richness, countering the effects of infection. However, we recognised a few limitations in this study, as it is a descriptive study where non-probability sampling was carried out with a relatively small sample size. Moreover, this study highlighted results at the phylum level but did not evaluate at the genus level. Also, we did not address the link between *Blastocystis* STs and gut microbiota modification, which could be a piece of crucial information to determine the alteration in the gut as different STs potentially can act as commensal or pathogen. Nevertheless, the role of the *Blastocystis* subtypes did not clearly explain in the gut as the hypothesis needs further investigations (27).

## Conclusion

To the best of our knowledge, this is the first study comparing the gut microbiota’s characteristics of Malaysian adults with T2DM and non-diabetic adults in the presence with and without *Blastocystis* utilising amplicon-based sequencing of 16S rRNA genes. This study provided insight into the most representative phyla behaviour in both T2DM and non-diabetic groups when *Blastocystis* is present, resulting in higher diversity and richness of the gut bacterial composition. A potential relationship between *Blastocystis* and gut microbiota should be elucidated as it could be associated with healthy intestinal microbiota or dysbiosis depending on many factors. Despite, animal models have highlighted some modifications specifically to this parasite, however, *Blastocystis* interaction towards gut composition changes remains unknown. Thus, further studies should emphasise on understanding the transmission mechanism and pathogenicity in humans with evaluation of the composition of the gut is dependent on a more precise assessment of *Blastocystis* diversity.

## Acknowledgements

The authors are very grateful to the Ministry of Higher Education, Malaysia, the research management centre at Universiti Sains Islam Malaysia for financing and providing help and storage resources. A special thanks also go to all the volunteers and medical staff from Pusat Perubatan Universiti Kebangsaan Malaysia (PPUKM) for their technical assistance in samples and data collection.

